# Characterization and functional interrogation of SARS-CoV-2 RNA interactome

**DOI:** 10.1101/2021.03.23.436611

**Authors:** Athéna Labeau, Alain Lefevre-Utile, Lucie Bonnet-Madin, Luc Fery-Simonian, Vassili Soumelis, Vincent Lotteau, Pierre-Olivier Vidalain, Ali Amara, Laurent Meertens

**Affiliations:** INSERM U944 CNRS 7212, Université de Paris, Institut de Recherche Saint-Louis, Hôpital Saint-Louis, 75010 Paris, France; INSERM U976, Université de Paris, Institut de Recherche Saint-Louis, Hôpital Saint-Louis, 75010 Paris, France; Centre International de Recherche en Infectiologie (CIRI), Inserm U1111, Université Claude Bernard Lyon 1, CNRS UMR5308, ENS de Lyon, 69007 Lyon, France

## Abstract

Severe acute respiratory syndrome coronavirus 2 (SARS-CoV-2) is the causative agent of COVID-19 pandemic, which has caused a devastating global health crisis. The emergence of highly transmissible novel viral strains that escape neutralizing responses emphasizes the urgent need to deepen our understanding of SARS-CoV-2 biology and to develop additional therapeutic strategies. Using a comprehensive identification of RNA binding proteins (RBP) by mass spectrometry (ChIRP-M/S) approach, we identified 142 high-confidence cellular factors that bind the SARS-CoV-2 viral genome during infection. By systematically knocking down their expression in a human lung epithelial cell line, we found that the majority of the RBPs identified in our study are proviral factors that regulate SARS-CoV-2 genome replication. We showed that some of these proteins represented drug targets of interest for inhibiting SARS-CoV-2 infection. In conclusion, this study provides a comprehensive view of the SARS-CoV-2 RNA interactome during infection and highlights candidates for host-centered antiviral therapies.

## Introduction

SARS-CoV-2, which belongs to the *Coronaviridae* family, has been identified as the causative agent of the ongoing pandemic of coronavirus disease 2019 (COVID-19) (Zhu et al., 2020). As of March 7, 2021, SARS-CoV-2 has spread Worldwide, with about 116 million confirmed cases and above 2.5 million deaths according to the WHO. Vaccination appears as one of the most expected health interventions with over 200 vaccine candidates under development as of December, 2020 (Mellet and Pepper, 2021) with those that received an emergency approval targeting the viral spike protein (Poland et al., 2020). However, the emergence of SARS-CoV-2 variants with mutations in the spike protein that reduces susceptibility to neutralizing antibodies (Choi et al., 2020; Kemp et al., 2021; Liu et al., 2021; McCarthy et al., 2021; Thomson et al., 2021), raises the concern of future variants escaping from natural or vaccine-induced neutralizing immunity. Furthermore, a recent study conducted in Manaus, Brazil, suggested a potential antibody wane at 8 months post-exposure (Sabino et al., 2021). Thus, there is an urgent need to improve our understanding of SARS-CoV-2 replication mechanisms to develop additional therapeutic strategies.

The genome of SARS-CoV-2 is composed of a single stranded positive RNA with a length of approximatively 30 kb (Zhou et al., 2020). In the cytoplasm of infected cells the viral RNA is translated by the host machinery in two overlapping replicase polyproteins that undergo subsequent viral-protease-mediated processing to generate 16 non-structural proteins (NSP) (de Wilde et al., 2018). These NSPs assemble and recruit poorly-characterized host cell factors to form the replication and transcription complex (RTC), which localizes in a virus-induced network composed of double-membrane vesicles (DMV)(Cortese et al., 2020; Eymieux et al., 2021). Within the RTC, the RNA-dependent-RNA polymerase (RdRp) complex uses the genomic RNA as a template to generate negative strand intermediates that serve for the synthesis of both progeny genomic RNA and subgenomic viral RNA (sgvRNA) (Snijder et al., 2016). SgvRNAs are then translated in 4 structural (S, E, M and N) and seven accessory proteins (ORF3a-9b) (Kim et al., 2020). Viral genomic RNA associates with the nucleocapsid (N) to form the ribonucleocapsid that drives, with the other structural proteins, the assembly of new viral particles, which bud in the lumen of the endoplasmic reticulum-Golgi intermediate compartment (ERGIC).

Like all viruses, SARS-CoV-2 heavily relies on cellular proteins to accomplish its infectious life cycle. This host dependency represents a potential “Achilles’ heel” that could be exploited to develop new host-centered approaches to treat SARS-CoV-2 infection. In this context, several strategies were used to decipher the molecular interactions occurring between the virus and host cell during infection. Affinity purification combined with mass spectrometry allowed the identification SARS-CoV-2 virus-host protein-protein interactions (PPI) (Gordon et al., 2020a, 2020b; Li et al., 2020) while CRISPR-Cas9 phenotypic screens have provided valuable information about the host genes and cellular pathways important for the SARS-CoV-2 life cycle (Daniloski et al., 2020; Hoffmann et al., 2021; Schneider et al., 2020; Wang et al., 2021; Wei et al., 2020). These studies have led to the identification of potential cellular targets for drug repurposing, such as the PI3K complex, the sigma-1 and −2 receptors as well as the cholesterol homeostasis (Daniloski et al., 2020; Gordon et al., 2020a; Hoffmann et al., 2021; Wang et al., 2021).

In this study, using a comprehensive identification of RNA-binding protein by mass spectrometry (ChIRP-M/S) approach, we provide a global map of the host proteins that associate with the SARS-CoV-2 genome during infection. By combining these proteomics data with gene silencing experiments, we identified a set of new SARS-CoV-2 host dependency factors that impact infection and could be targetable for antiviral therapies.

## Results and Discussion

### Characterization of the SARS-CoV-2 RNA interactome in human cells

To characterize the host factors that interact with SARS-CoV-2 genome during infection, we performed a comprehensive identification of RNA-binding proteins by mass spectrometry approach (ChIRP-M/S) (Figure 1A). Originally developed to study endogenous ribonucleoprotein (RNP) complexes, this technique has been recently adapted to investigate viral RNA-host protein interactions (Ooi et al., 2019). Briefly, we challenged 293T cells stably expressing the human ACE2 receptor (293T-ACE2) with a primary clinical SARS-CoV-2 220/95 strain (EPI_ISL_469284) isolated by our laboratory in March 2020 (Onodi et al., 2021). Forty-eight hours later, SARS-CoV-2 infected and uninfected cells were treated with formaldehyde to crosslink RNA with RBPs in order to preserve RNP complexes. To specifically capture SARS-CoV-2 RNA, we designed 129 biotinylated antisense probes (Table S1) complementary of the fulllength viral RNA, and subsequently used streptavidin beads to pulldown the RNPs. Co-immunoprecipitated proteins were separated by SDS-PAGE, visualized by silver-staining (Figure 1B), and subjected to mass spectrometry (MS) analysis (Table S2). As expected, the SARS-CoV-2 N protein, which is the main viral RBP, was the most enriched protein in our study (Figure 1B, C). The structural M and S proteins, the replicase polyprotein pp1ab and the accessory protein ORF9b were also significantly and reproducibly enriched in our experiments (Figure 1C, 1D). Of the 16 NSPs released from pp1ab, 9 were enriched in our ChIRP-M/S (Figure 1D) and are part of the coronavirus RTC (V’kovski et al., 2019), where NSP2-NSP16 play essential functions involved in RNA replication (V’kovski et al., 2021). NSP1 is known to interact with 40S subunit of the ribosome to inhibit cellular translation, and to bind the viral 5’ untranslated region (UTR) to promote vRNA translation (Schubert et al., 2020). The structural protein M and S associate with the viral RNA during virus assembly and budding. Finally, ORF9b has be shown to localize at the mitochondria membrane and antagonize the RIG-I-MAVS antiviral signaling through its association with TOM70 (Gordon et al., 2020b; Jiang et al., 2020). Together, these data provide strong evidences that our ChIRP-M/S approach reproductively and specifically pulldown viral RNP complexes involved in different steps of SARS-CoV-2 life cycle, from replication/transcription to particle assembly.

**Figure 1.**
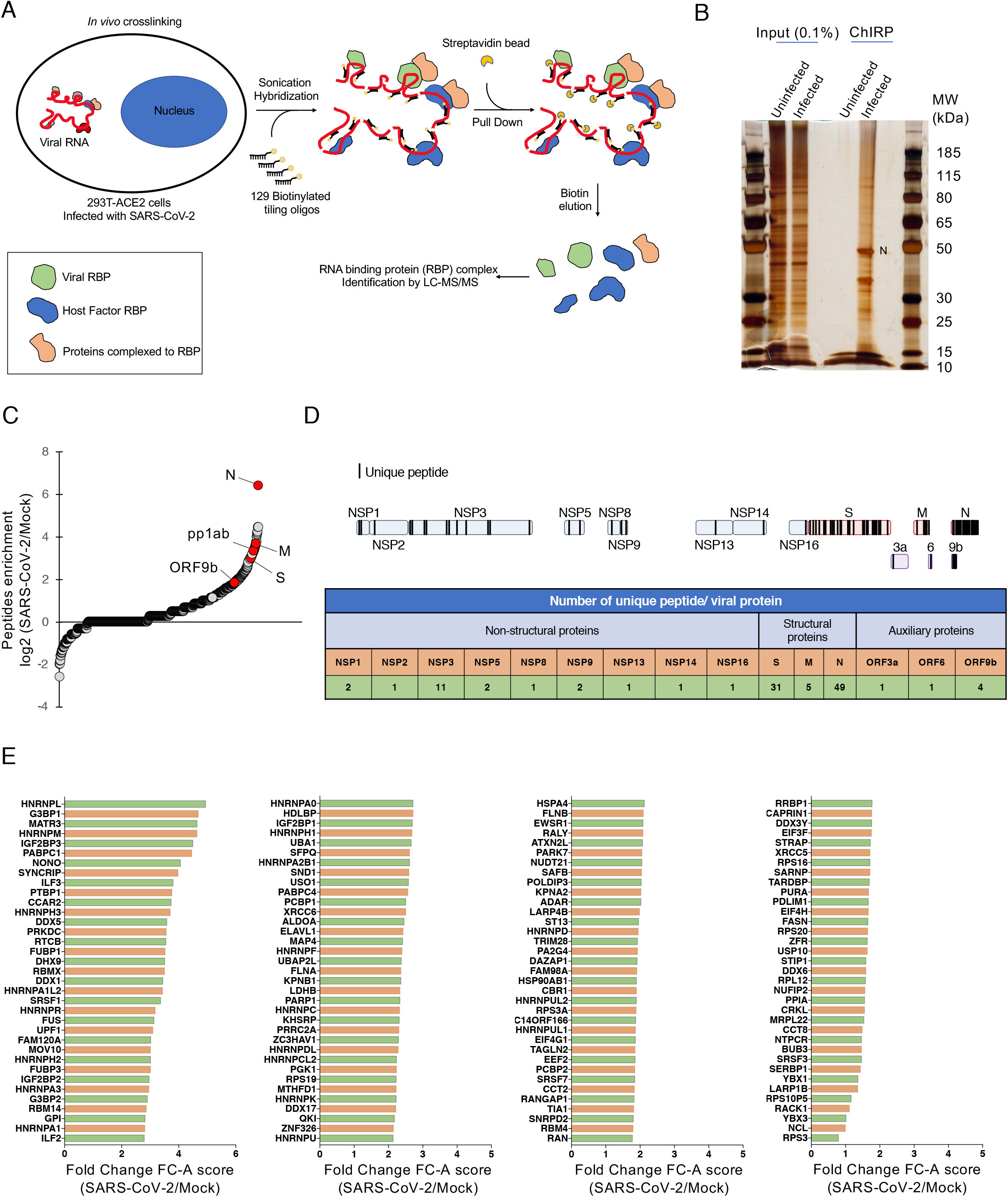
Purification of the SARS-CoV-2 RNA interactome. **(A)** Schematic illustrating the ChIRP-M/S approach used to identify host factors bound to SARS-CoV-2 RNA. **(B)** Proteins enriched by ChIRP from uninfected and SARS-CoV-2-infected cells were resolved by SDS-PAGE and visualized by silver staining. **(C)** Quantification of the peptides enrichment in SARS-CoV-2 infected cells over uninfected cells (n=5 independent replicates). Viral proteins are marked by a red circle. **(D)** (top) Schematic representing unique peptides distribution along the enriched viral proteins. (bottom) Table showing the number of unique peptides for each viral protein. Data from one representative replicate were shown. **(E)** Fold change (SAINTexpress FC-A score) of the 140 high-confident host proteins identified in our SARS-CoV-2 ChIRP-M/S. (Proteins Histone H3.2 and H2B type 1-H were not represented).

### A view of the SARS-CoV-2 RNA-host protein interactome

Using SAINTexpress scoring algorithms, we identified 142 high-confident human proteins that reproductively associated with the SARS-CoV-2 RNA (SAINT score ?0.79; Figure 1E, Table S3). Among them, 89 % (127 out of 142) were annotated by the ontology term “RNA binding” (GO: 0003723), suggesting a direct interaction with the viral genome. Next, we compared our RNA interactome with recently published data sets obtained by RNA-centric (RAP-M/S and ChIRP-M/S) approaches (Flynn et al., 2021; Lee et al., 2020; Schmidt et al., 2020). We found that 24% to 38% (Schmidt *et al*.: 35/144; Flynn *et al*.: 77/228 and *Lee et al*.: 74/193) of the SARS-CoV-2 RNA host factors identified in these studies were also enriched in our ChIRP-M/S interactome (Table S4). Interestingly, we highlighted 58 common hits which defined a reproducible set of cellular SARS-CoV-2 RNA RBP (Figure S1). These include multiple RBPs involved in nucleic acid metabolism including splicing, mRNA stability and transcriptional regulation such as several heterogenous nuclear ribonucleoproteins (hnRNPs) (Geuens et al., 2016), the serine/arginine-rich splicing factor (SRSF) (Shepard and Hertel, 2009), poly(A) binding protein (PABP) (Mangus et al., 2003) and the Insulin-like growth factor 2 mRNA binding protein (IGF2BP) (Bell et al., 2013). We also found several components of the small 40S ribosome subunit (RPS3, RPS3A, RPS16, RPS19 and RPS20), components of the translation initiation complex (EIF4G1, EIF3F and EIF4H) and translation elongation factor (EEF2). Surprisingly, we did not identify EIF4E, the CAP binding protein, suggesting that SARS-CoV-2 might use an alternate CAP-dependent EIF4E-independent translation mechanism (de la Parra et al., 2018). Chaperone proteins (HSP90AB1, CCT2 and CCT8) were also significantly enriched and are likely involved in viral protein folding. Characteristic RBPs involved in ribonucleoprotein granules such as paraspeckles (PS) (RBM14, NONO, MATR3, SFPQ) and stress granules (SG) (G3BP1, G3BP2, TIA1, CAPRIN1, ATXN2L) (Nakagawa et al., 2018; Yang et al., 2020) were among the highest-scoring candidates. Many PS-interacting proteins were also highly represented in our ChIRP-M/S data set (DAZAP1, ESWR1, FAM98A, FUS, RBMX, UBAP2L, as well as hnRNPA1, A1L2, H1, H3, K, R, UL1) (Naganuma et al., 2012).

Functional analyses of the RBPs revealed more than 150 statistically enriched GO terms, which cluster into 23 functional groups (Figure 2A), and are also found in the three other SARS-CoV-2 RNA interactome studies (Figure 2B). Among them, we found host molecules that participate in RNA metabolic process, translation, RNA stabilization, ribonucleoprotein complex assembly, viral process and innate immune response (Figure 2A, 2C). Based on its predominant nuclear localization and its role in mRNA maturation, the significant enrichment of RNA splicing GO term (p=3.55×10^-43^) was unexpected. Pre-mRNA splicing is critical for mRNA nucleo-cytoplasmic export, thereby contributing to the regulation of gene expression (Reed and Hurt, 2002). It is tempting to speculate that SARS-CoV-2 RNA may sequester host RBP involved in splicing in order to hamper cellular mRNA nuclear export, thus favoring the cytoplasmic translation of viral RNAs. This hypothesis is consistent with recent findings showing that SARS-CoV-2 NPS16 disrupts mRNA splicing to antagonize innate immunity at a post-transcriptional level (Banerjee et al., 2020).

**Figure 2.**
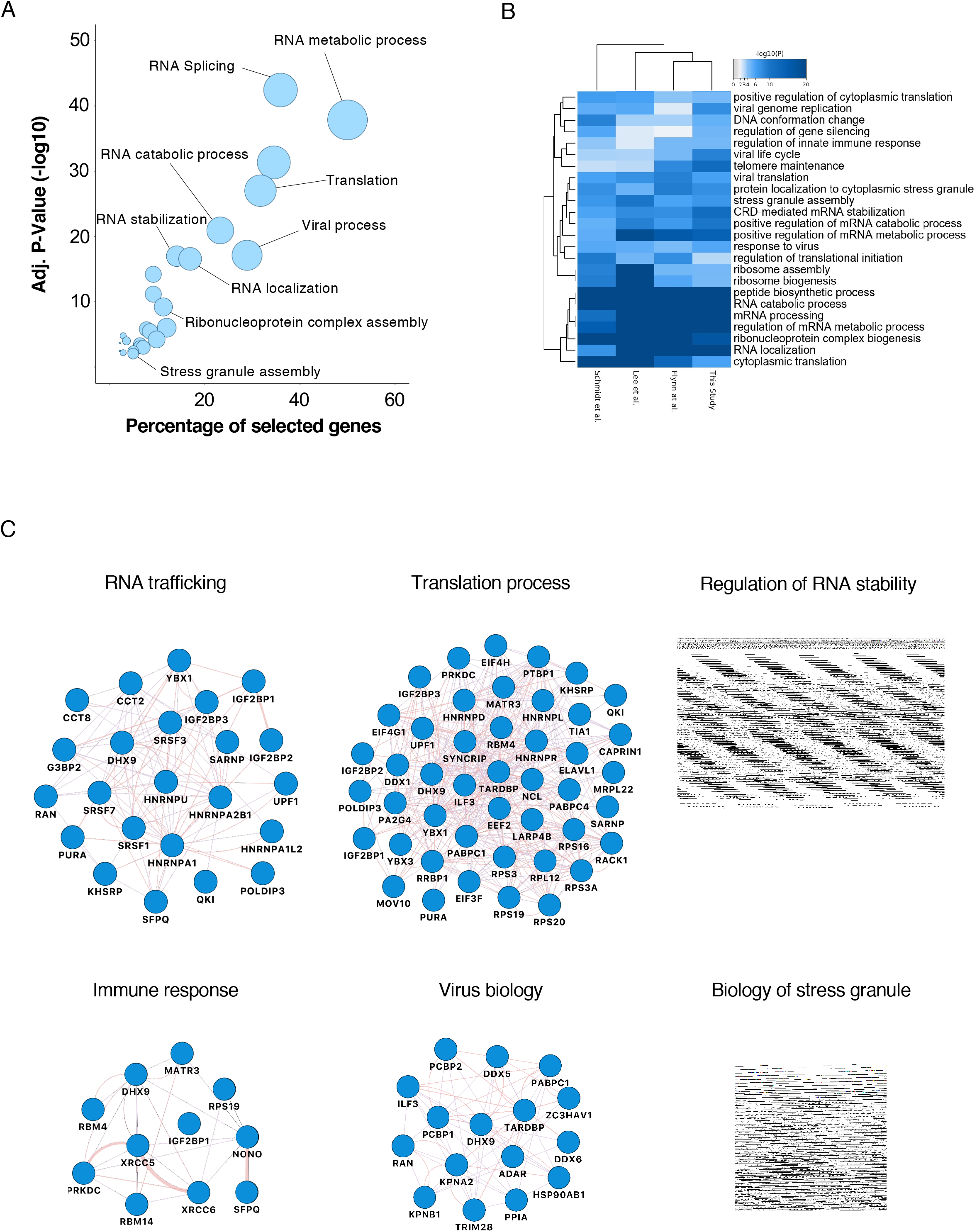
Biological analysis of the SARS-CoV-2 RNA interactome. **(A)** GO enrichment analysis of SARS-CoV-2 RNA interactome proteins. Circles represent enriched function for an annotated ontology term and circle size correspond to the number of enriched proteins within that term. **(B)** Heatmap of the top-ranked GO biological processes enriched across all SARS-CoV-2 RNA interactomes **(C)** Selected biological process networks of SARS-CoV-2 interactome proteins.

### Overlapping of the ChIRP interactome with known coronavirus-host interactions identifies host factors involved in central carbon metabolism as a new target of SARS-CoV-2

To gain insights on the interactions between host factors and viral proteins we compared our SARS-CoV-2 interactome with a reference coronavirus interactome, which corresponds to a meta-analysis of the viral-host protein-protein interactions of 13 coronaviruses collected built upon 112 publications (Perrin-Cocon et al., 2020). Of the 142 high-confidence hits identified by our ChIRP approach, 80 overlapped with the 1,140 host proteins (56%) of the reference coronavirus interactome. As illustrated in Figure 3A, these host proteins captured by our ChIRP assay are known interactors of one or several coronaviruses viral proteins. Interestingly, a majority of these proteins (79%) were previously identified as N protein-binding partners. Shared interactors with S, NSP1, NSP2, NSP9, NSP12, NSP13, NSP14, NSP15, ORF6 and ORF9b were also identified (Figure 3A). In our interaction network, the ATP-dependent RNA helicase DDX1 was the most connected factor interacting with both N and NSP14 from both TGEV and PEDV, NSP14 from SARS-CoV-1 and NSP2 from MHV. Altogether, this supports a key role of DDX1 in SARS-CoV-2 replication cycle, as previously reported for other coronaviruses (Wu et al., 2014; Xu et al., 2010).

**Figure 3.**
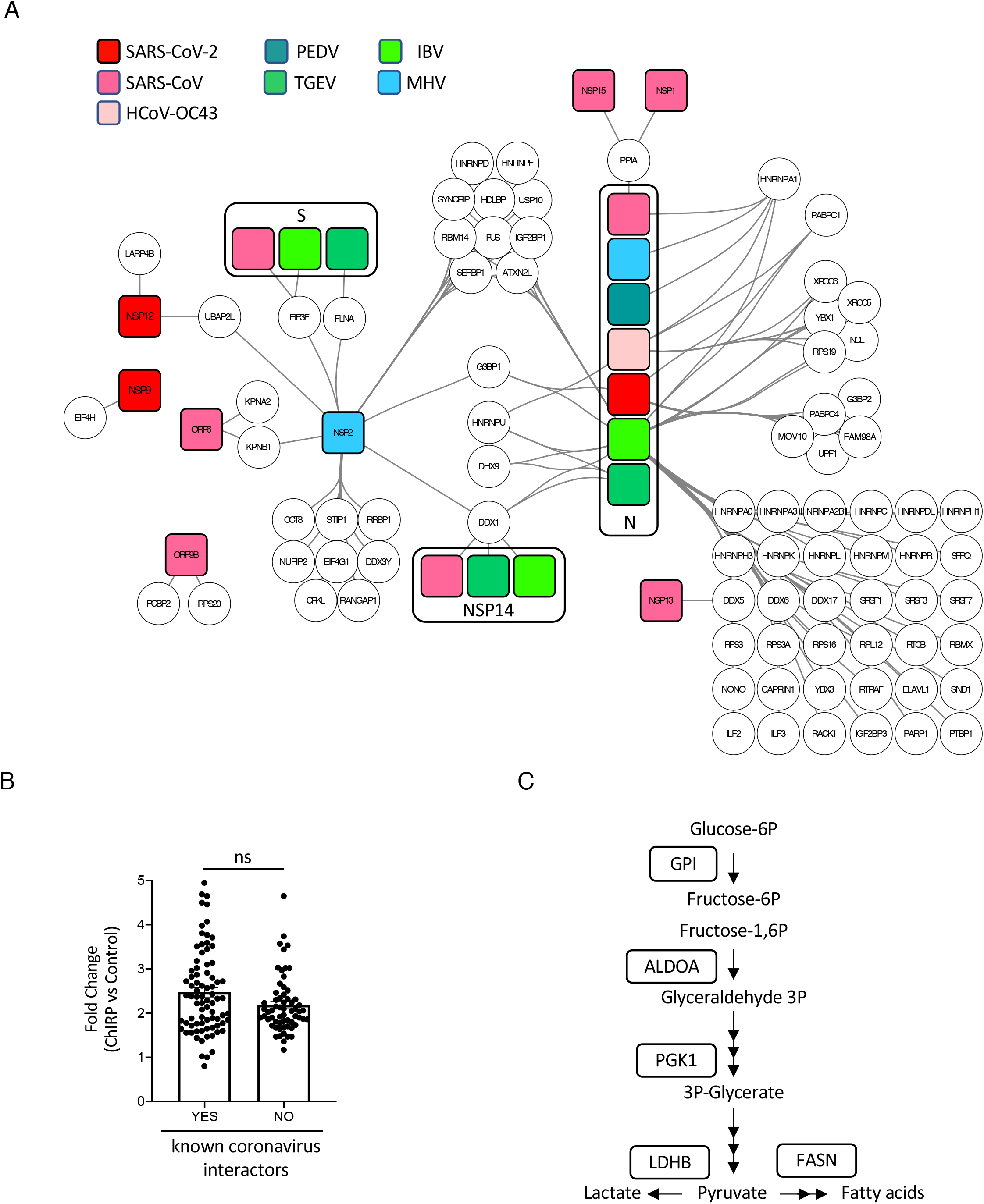
Overlapping of our ChIRP interactome with known coronavirus-host interactions identifies the central carbon metabolism as SARS-CoV-2 target. **(A)** Map showing interactions between the cellular proteins of our ChIRP dataset and viral proteins of SARS-CoV-2 or other coronaviruses. This analysis was performed using the reference coronavirus interactome previously published (Perrin-Cocon et al., 2020). Round circles correspond to host proteins. Squares correspond to viral proteins. **(B)** Fold change scores of cellular proteins that are either known coronavirus targets or new interactors identified for the first time by the ChIRP study. Fold change scores (FC-A) were calculated by Saint-Express from the average mean of spectral counts. No significant differences were observed between the two distributions (p-value>0.05; Mann Whitney test). **(C)** Simplified metabolic pathways showing functional connections between five enzymes identified in our ChIRP analysis and targeted by SARS-CoV-2.

We then analyzed the subset of cellular proteins that did not overlap with the reference coronavirus interactome, which essentially corresponds to novel SARS-CoV-2 and coronavirus interactors. First, we compared the fold enrichment in the ChIRP experiments for the 62 new and 80 known interactors and observed a similar distribution, indicating that the two subsets were qualitatively equivalent and correspond to robust interactions (Figure 3B). Next, we investigated distribution of biological processes covered by the subset of new interactors, and found a distribution of the functional annotations similar to what we observed with the full data set. However, eight proteins (ALDOA, CBR1, FASN, GPI, LDHB, MTHFD1, NTPCR and PGK1) belonged to metabolic pathways according to KEGG database. Interestingly, five proteins (GPI, ALDOA, PGK1, LDHB, FASN) were involved in glycolysis, lactagenesis and lipogenesis which are tightly connected pathways (Figure 3C). These data are consistent with recent a study showing that glycolysis and lactate pathways sustained SARS-CoV-2 replication (Codo et al., 2020). Furthermore, fatty acid synthesis is necessary for the formation of lipid droplets (LD) which are required for SARS-CoV-2 viral replication (Dias et al., 2020). Together, these data suggest that SARS-CoV-2 targets the central carbon metabolism to accomplish its life cycle.

### Functional interrogation of the SARS-CoV-2 RBP identified in our ChIRP-M/S approach

To further pinpoint the function of the identified RBPs in SARS-CoV-2 infection, we silenced their expression by RNA interference (RNAi) and determined consequences on virus infection. We designed a library of siRNA pool targeting 138 RBPs identified in our study and performed a loss-of-function screen in the human lung epithelial cell line A549 stably expressing ACE2. SiRNA targeting the cathepsin L protease (CTSL) and the vacuolar ATPase subunit ATP6V1B2, two host molecules important for SARS-CoV-2 viral entry (Daniloski et al., 2020), were used as positive controls, whilst a non-targeting siRNA pool (siNT) was used as negative control. SiRNA transfected cells were challenged with SARS-CoV-2 for 24 hours and viral infection was assessed either by quantifying the intracellular accumulation of the N protein by flow cytometry (viral replication) or the viral RNA released in the supernatant of infected cells by RT-qPCR (release of viral particles) (Figure 4A). All readouts were normalized to the siNT-transfected cells. Transfected cells were also monitored for cell viability, and 7% (9/138) of the siRNA affecting cell viability were removed from the analysis. As expected, transfection of siRNA targeting CTSL and ATP6V1B2, resulted in a strong decrease of infection in both assays (Figure 4B-C). RT-qPCR experiments revealed that 45 siRNAs reduced by at least 50% viral particles production whereas 4 siRNAs increased it by 200 % (Figure 4B). Flow cytometry studies showed that 21 siRNAs decreased the percent of infected cells by more than 50 %, while 12 enhanced it by 150% (Figure 4C). By intersecting the data from both assays, we identified 25 RBPs that impacted viral infection. Interestingly, most of them (19/25) are host dependency factors (HDF) and few (6/25) act as host restriction factors (HRF) (Figure 4D). This contrasts with the data recently published by Flynn et al. showing that the majority of SARS-CoV-2 RBP factors identified in their study has an antiviral function (Flynn et al., 2021). We observed a high correlation coefficient (r=0.742) between our RT-qPCR and flow cytometry data set indicating that most of the identified RBPs are required for viral replication (Figure 4D), and only very few act at a late step (assembly/particle egress) of the SARS-CoV-2 life cycle (Figure 4D, green circles).

**Figure 4.**
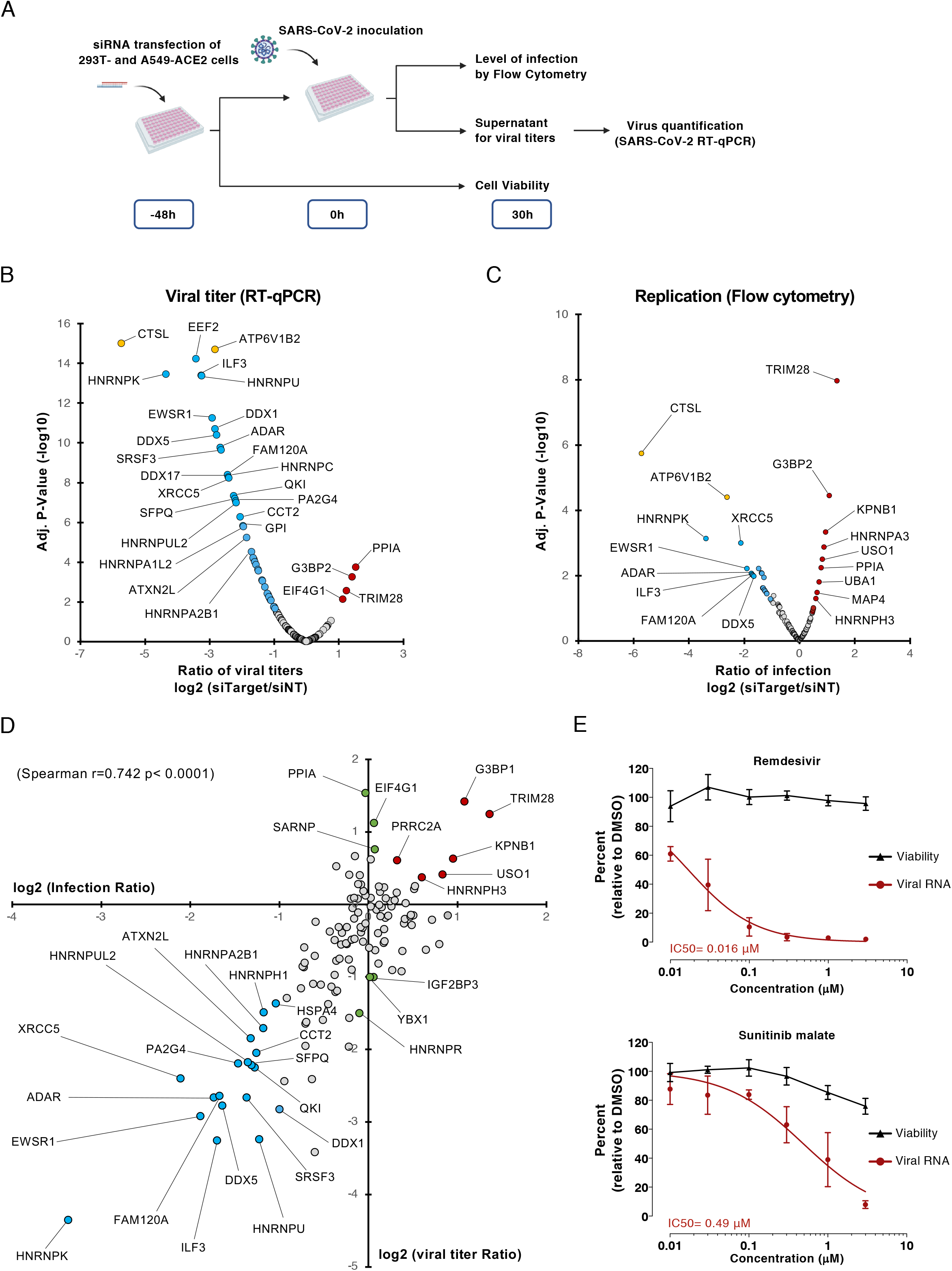
Functional interrogation of the SARS-CoV-2 RNA interactome and compounds screening. **(A)** Schematic illustrating the loss-of-function screen procedure. **(B and C)** A549-ACE2 cells were transfected with an arrayed siRNA library and challenged with SARS-CoV-2 (MOI 0.05) for 24h hours. **(B)** Yield of viral particles released in the supernatant of infected cells was quantified by RT-qPCR and normalized to the siNT-transfected cells. **(C)** Viral replication was assessed by flow cytometry using anti-N protein mAb, and normalized to the siNT-transfected cells. Data shown are means of two independent experiments. Adjusted p-values were calculated by one-way ANOVA with Benjamini and Hochberg correction. Host dependency factors are marked in blue and host restriction factors are marked in red. Positive controls (CTSL and ATP6V1B2) are highlighted in yellow. **(D)** Intersection of the data obtained from N protein quantification by flow cytometry and virus release in supernatant of infected cells by RT-qPCR. Data shown are means of two independent experiments. Host dependency factors are marked in blue and host restriction factors are marked in red. **(E)** A549-ACE2 were infected with SARS-CoV-2 (MOI 0.05) in continuous presence of increased concentrations of remdesivir or sunitinib malate. Virus released in supernatant was quantified 24 hpi by RT-qPCR (red lane). Cell viability was assessed in parallel (black lane). Data shown are mean +/- SD of three independent experiments in duplicate.

Interestingly, several of HDF such as ILF3, HNRNPU, HNRNPK, FAM120A and QKI were predicted to bind the 5’- or 3’-UTR of SARS-CoV-2 RNA (Sun et al., 2021). Others, including SFPQ, DDX5, EWSR1, HNRNPK, ADAR, PA2G4 were previously reported to promote replication of other RNA viruses (Garcia-Moreno et al., 2019; Gélinas et al., 2011; Landeras-Bueno et al., 2011; Li et al., 2013; Oakland et al., 2013; Thompson et al., 2020; Zhou et al., 2019a), supporting the role of these RBPs in SARS-CoV-2 replication. Interestingly, ILF3 was shown to participate to antiviral type I interferon response by promoting the expression of *IFNB1* and interferon-stimulated gene (ISG) during host translational shutoff (Watson et al., 2020). QKI was shown to repress host interferon response by downregulating MAVS expression at a post-transcriptional level (Liao et al., 2020). This suggests that SARS-CoV2 has evolved several mechanisms to dampen the type I interferon response by targeting specific RBPs.

Next, we looked at the DrugBank, ChEMBL, DGIdb (Drug-gene interaction database) and PanDrugs database for drugs that target our host RBP. We identified camptothecin (XRCC5), cerulenin (FASN), dabigatran (HNRNPC), geldanamycin (HSP90AB1), phenethyl isothiocyanate (HNRNPK), sunitinib malate (EWSR1) and artemisinin (an antimalarial interacting with multiple RBPs) as potential SARS-CoV-2 antiviral candidates. To test whether these compounds impact infection, A549-ACE2 cells were incubated with the compounds at a concentration of 10 and 1 μM and then challenged with SARS-CoV-2. We used equivalent dilution of DMSO as negative control and similar concentration of remdesivir, an inhibitor SARS-CoV-2 replication (Wang et al., 2020) as positive control. We monitored the release of progeny viruses in the supernatant as well as cell viability 24 hours post-infection. Of all the compounds tested, we observed a significant inhibitory effect for phenethyl isothiocyanate (40 % inhibition) and sunitinib malate (80 % inhibition) at a concentration of 1 μM without toxicity (Figure S2A). Sunitinib malate is a promising therapeutic for the treatment of patient suffering from sarcoma such as DSRCT (Desmoplastic Small Round Cell Tumor) or EMC which expressed an EWSR1-WT1 or EWSR1-NR4A3 fusion, respectively (Mello et al., 2021; Stacchiotti et al., 2014). Sunitinib malate inhibited the release of SARS-CoV-2 particles in infected cells in dose-response manner with an IC50 of 0.49 μM (Figure 4E). Sunitinib malate inhibits also the N protein production in infected cells, consistent with a role of EWSR1 in genome replication (Figure S3B). Similar results were obtained in 293T-ACE2 cells (Figure S3C).

In conclusion, our data provides a landscape of functional interactions that the SARS-CoV-2 RNA genome establishes with the host cell during infection. We provide several evidences that SARS-CoV-2 exploit several cellular RBP to facilitate viral replication and highlight host molecules that could be targeted for antiviral intervention.

## ACKNOWLEDGMENTS

This works was supported by French Government’s Investissement d’Avenir program, Laboratoire d’Excellence “Integrative Biology of Emerging Infectious Diseases” (grant n°ANR-10-LABX-62-IBEID), the Fondation pour la Recherche Medicale (grant FRM - EQU202003010193 and FDM201806006187), the Agence Nationale de la Recherche (ANR) (ANR-20-COVI-000 project IDISCOVR and ANR-20-CO11-0004 project FISHBP) and University of Paris (Plan de Soutien Covid-19: RACPL20FIR01-COVID-SOUL). Laurent Meertens and Ali Amara dedicate this work to the memory of Professor Jean-Louis Virelizier (Unité d Immunologie Virale, Institut Pasteur, Paris) and Professor Renaud Mahieux (Ecole Normale Supérieure, Lyon, France), who left us during the SARS-CoV-2 epidemic. The authors thank Alessia Zamborlini for critical reading of the manuscript.

## AUTHOR CONTRIBUTION

Conceptualization, L.M. and A.A.; L.M. supervised the project. L.M. performed the ChIRP experiments and analyzed the data with the help of A.L and P.O; A.L-U.; P.O. and V.S. performed the bioinformatic analysis; L.B-M and L.M produced the SARS-CoV-2 virus stocks used in this study. L.F-S participated in the ChIRP experiments. P.O.V and V.L. analyzed the viral protein-RBP protein interaction; A.L and L.M. performed the siRNA screen; A.L. screened the compounds; L.M., P.O.V., A.L. and A.A. wrote the manuscript with the inputs from all authors; Funding Acquisition, A.A and L.M.

## DECLARATION OF INTEREST

The authors declare no competing interest.

## Tables titles

**Table S1.** Sequences of the 129 biotinylated ChIRP probes. Related to figure 1.

**Table S2.** Datasets of the mass spectrometry analyses. Related to figure 1.

**Table S3.** SAINTexpress analyses of the ChIRP-M/S dataset. Related to figure 1.

**Table S4.** List of RBP identified in ChIRP-M/S that overlap with other SARS-CoV-2 interactome. Related to figure 1.

## Supplementary Figure Legends

**Figure S1.**
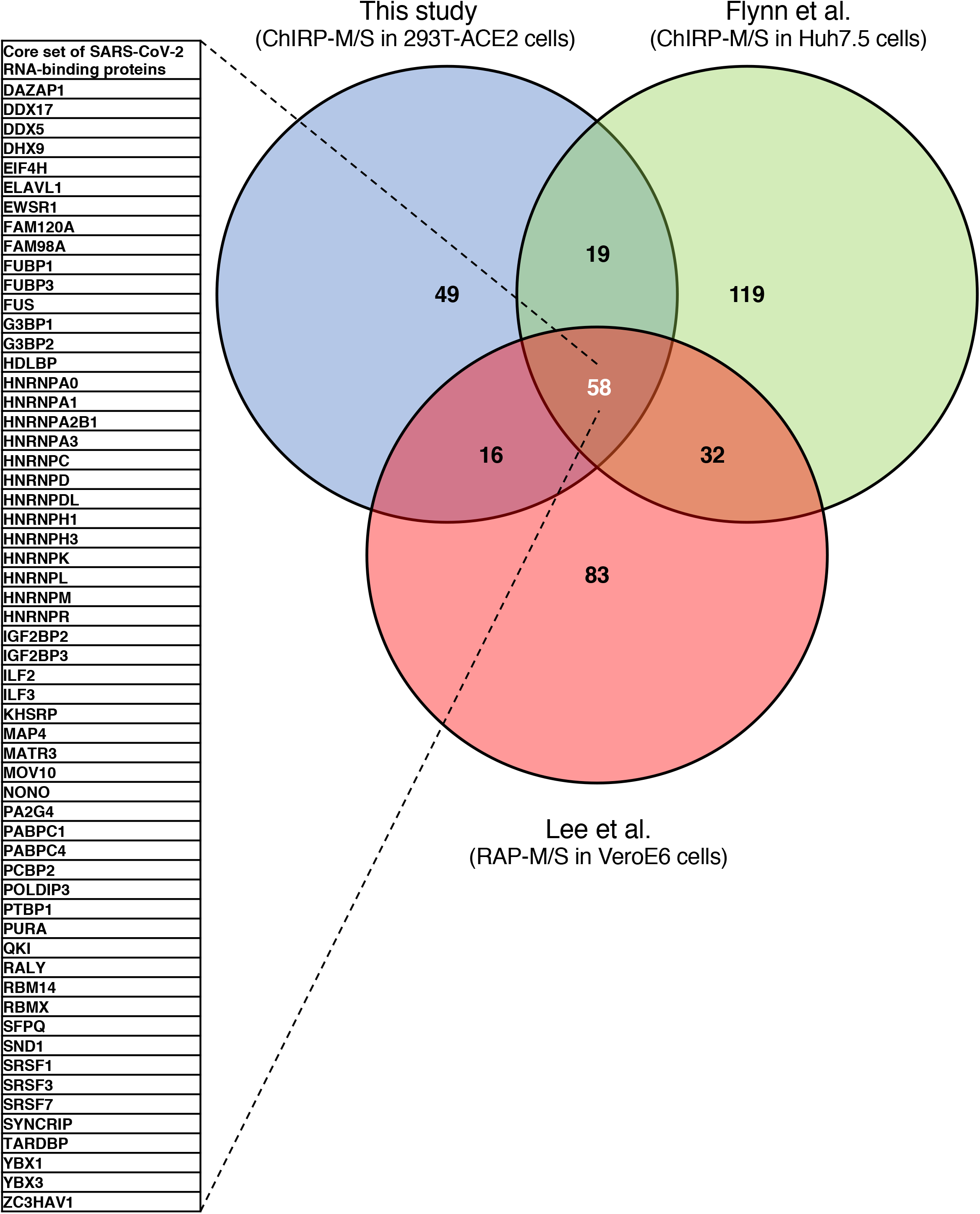
Intersection of SARS-CoV-2 interactomes identified a set of core SARS-CoV-2 RNA RBP. Related to Figure 1. Venn Diagram comparing the significantly enriched RBP from SARS-CoV-2 RNA interactomes.

**Figure S2.**
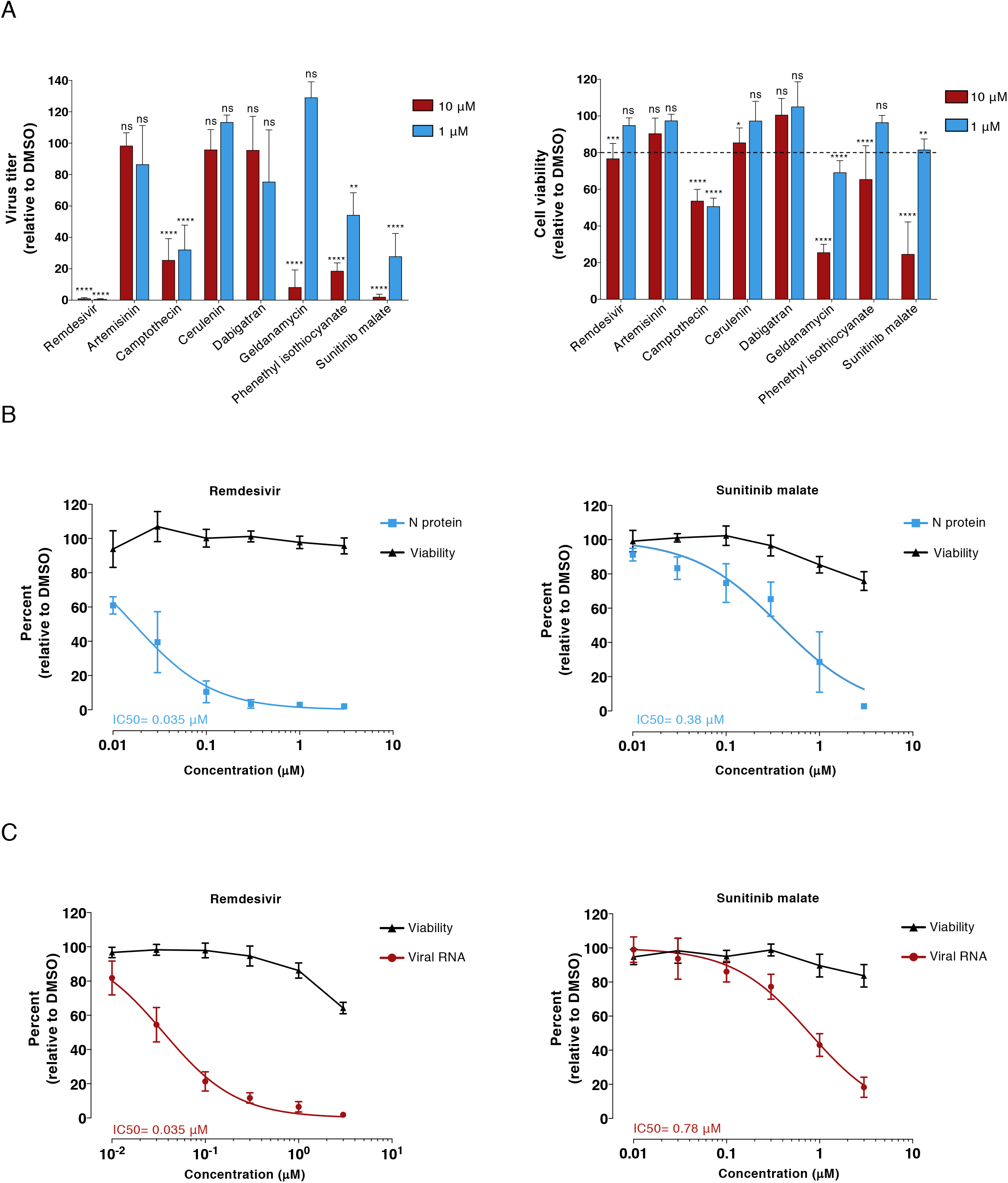
Screening of compounds with antiviral activity targeting SARS-CoV-2 host RBP. Related to Figure 4. **(A)** A549-ACE2 were infected with SARS-CoV-2 (MOI 0.05) in continuous presence of compounds (10 and 1 μM). Virus released in supernatant was quantified 24 hpi by RT-qPCR (top panel). Cell viability was assessed in parallel (bottom panel). Data shown are mean +/- SD of three independent experiments in duplicate. Significance was calculated using two-way ANOVA statistical test with Dunnett’s multiple comparisons test. (ns not significant, ** p< 0.01; *** p< 0.001; **** p<0.0001). (B) A549-ACE2 were infected with SARS-CoV-2 (MOI 0.05) in continuous presence of increased concentrations of remdesivir or sunitinib malate. Percentage of infected cells was quantified 24 hpi by flow cytometry using anti-N mAb (blue lane). Cell viability was assessed in parallel (black lane). Data shown are mean +/- SD of three independent experiments in duplicate. (C) 293T-ACE2 were infected with SARS-CoV-2 (MOI 0.05) in continuous presence of increased concentrations of remdesivir or sunitinib malate. Virus released in supernatant was quantified 24 hpi by RT-qPCR (red lane). Cell viability was assessed in parallel (black lane). Data shown are mean +/- SD of three independent experiments in duplicate.

## EXPERIMENTAL METHODS

### Cell lines

Lenti-X 293T cells (human embryonic kidney cells), A549 cells (Human alveolar basal epithelial carcinoma), and Vero E6 cells (African green monkey kidney cells) were maintained in Dulbecco Modified Eagle Medium (DMEM; Invitrogen Life Technologies) supplemented with 10% heat-inactivated fetal bovine serum (FBS) and 1% penicillin/streptomycin (P/S) and 1% GlutaMAX (Life Technologies). All cell lines were periodically tested negative for mycoplasma contamination prior to use in experiments.

### Virus preparation and titration

SARS-CoV-2 strain 220_95 (EPI_ISL_469284) was isolated from nasopharyngeal swab specimens collected at Service de Virologie (Hospital Saint Louis, Paris) (Onodi et al., 2021). Virus was propagated on Vero E6 in DMEM-2% (DMEM supplemented with 2% FBS, 1% P/S, 1% GlutaMAX, and 25 mM Hepes). Viruses were passaged three times before being used for experiments. For the last passage, viruses were purified through a 20% sucrose cushion by ultracentrifugation at 80,000xg for 2 hours at 4°C. Pellets were resuspended in TNE1X pH7.4 (Tris 50 mM, NaCl 100mM, EDTA 0.5 mM), aliquoted and stored at −80°C.

Viruses titer was ascertained by plaque assays in Vero E6 cells and expressed as PFU per ml. Cells were incubated for one hour with 10-fold serial dilution of viral stocks. The inoculum was then replaced with Avicel 2.4% mixed at equal volume with DMEM supplemented with 4% FBS, 2% Glutamax, 50mM MgCl_2_, 0.225 % of NaHCO_3_, and incubated 3 days at 37°C. Then, Vero E6 cells were washed twice with phosphate-buffered saline (PBS) and stained with an aqueous solution containing 1% formaldehyde, 0,5% crystal violet, and 0,89% sodium chloride, for 15 min at room temperature. Vero E6 cells were then washed with water to visualize lysis plaques.

### Generation of cell lines stably expressing ACE2

The human ACE2 coding DNA sequence (CDS) was amplified by RT-PCR from pLenti-ACE2 plasmid (a gift from Olivier Schwartz, Virus and Immunity, Institut Pasteur) using the ACE2 forward and reverse primers. Amplification product was cloned into pLVX-IRES-ZsGreen1 vector using XhoI/NotI restriction enzymes. The ACE2 protein-expressing lentiviral vectors pseudotyped with vesicular stomatitis virus glycoprotein G (VSV-G) were generated by transfecting lenti-X 293T (5×10^6^) cells with pLVX-ACE2-ZsGreen, psPAX2, and pVSV-G (4:3:1 ratio) using calcium phosphate transfection (Promega) following the manufacturer’s instructions. Supernatants were harvested 48 hours after transfection, cleared by centrifugation, and filtered. Lentiviral vectors were purified as describe above (see ‘virus preparation and titration’ section). 293T and A549 cells (2×10^5^) were seeded in 6 well plate and transduced by spinofection at 1,000xg for 2 hours at 33°C. Cell populations were sorted based on GFP expression with a BD FACSAria II (Becton Dickinson) with FACSDiva 6.1.2 software (Becton Dickinson).

### Flow cytometry analysis of ACE2 cell surface expression

Cells were detached with 2.5 mM EDTA in PBS and incubated with anti-human ACE2 mAb (5 mg/ml) in 100 ml of PBS with 0.02% NaN3 and 5% FBS for 1 hour at 4°C. Following the primary staining, cells were washed and stained with incubated with the Alexa 647-conjugated goat anti-mouse IgG (Jackson ImmunoResearch) for 30 min at 4°C. Acquisition was performed on an Attune NxT Flow Cytometer (Thermo Fisher Scientific) and analysis was done by using FlowJo software (Tree Star)

### Comprehensive identification of RNA binding proteins by mass spectrometry

ChIRP-M/S was performed as previously described by Ooi and colleagues (Ooi et al., 2019). SARS-CoV-2 probes were designed using the online tool (https://www.biosearchtech.com/stellaris-designer), with a repeat masking setting of 5 and even coverage of the whole viral genome (1 oligos every 200 bp). Oligos were synthetized with 3’ biotin-TEG modification at Eurofins Genomics. The 129 oligos were resuspended at 100 μM and mixed at equal volume to prepare the probes pool. 293T cells stably expressing ACE2 were seeded at 9×10^6^ cells per T150 flask (3 flasks per condition) and inoculated with SARS-CoV-2 at a multiplicity of infection (MOI) of 0.001 or mock-treated. The media was removed 48 hours later and the cells were rinsed once with PBS, trypsinized and pooled for each condition. Cells were centrifugated at 1,500 rpm for 5 min and pellets were washed twice with PBS. The cells were resuspended with 35 ml of PBS containing 3% formaldehyde and incubated under agitation for 30 min at room temperature (RT). Of note, all the buffers used throughout the ChIRP experiments were freshly prepared. Cross-linking was terminated by adding glycine to a final concentration of 0.125 mM for 5 min at RT. Cross-linked cells were centrifugated at 1,000xg for 5 min, washed once with PBS, weighted, then snap frozen and kept at −80°C until used. Cells were resuspended in 1 ml of lysis buffer (50 mM Tris-HCl, pH7.0, 10 mM EDTA, 1% SDS supplemented with SUPERase-IN and protease inhibitor) per 100 mg of cell pellet. The lysates were sonicated using a Diagenode Bioruptor. At this stage, 10 μl of lysates were kept as sample inputs. Lysates were precleared by adding 30 μl of washed MyOne C1 beads per ml of lysate and rocked for 30 min at 37 °C. Beads were removed from the lysates using a magnetic stand and 2 ml of ChIRP hybridization buffer (750 mM NaCl, 1% SDS, 50 mM Tris-HCl pH 7.0, 1 mM EDTA, 15% formamide supplemented with 1 mM PMSF, protease inhibitor and SUPERase-in) and 1.5 μl of probes pool were added for every ml of lysate. After mixing, hybridization took place for 16 hours at 37 °C under rotation. Then, 100 μl of washed MyOne C1 beads per microliter of probes pool were added to each sample and incubated for 45 min at 37°C. Magnetic beads were collected on a magnetic stand and washed 5 times with 1 ml of washing buffer (2x SSC, 0.5% SDS, add 1 mM PMSF) for 5 min at 37°C. To eluate the enriched ribonucleoprotein complexes, the beads were incubated with 300 μl of biotin elution buffer (12.5 mM D-biotin, 7.5mM Hepes pH 7.5, 75 mM NaCl, 1.5 mM EDTA, 0.15 % SDS, 0.075% sarkosyl and 0.02% Na-Doexycholate) for 20 min at RT under rotation and then 15 min at 65 °C with shaking. Eluate was collected in a fresh tube and the beads subject to elution again. The two eluents were pooled (600 μl) and 150 μl of trichloroacetic acid (25% of the total eluate volume) was added, mixed vigorously and let overnight at 4°C for precipitation. The next day, the samples were centrifugated at 20,000 g for 45 min at 4°C. Pellets were washed once with ice-cold acetone and spun at 16,000 rpm for 5 min at 4°C. The acetone supernatant was removed, the tubes centrifuged briefly to carefully remove the residual acetone and the pellets were left to air-dry on the bench top. The pellets were solubilized in 30 μl of Laemmli 2X buffer with 20 mM DTT and boiled at 95 °C for 30 min with a periodic mixing (every 5 min) to reverse the cross-linking.

One-tenth of the protein samples and their corresponding inputs (pre-diluted 1/10) were size-separated on a bolt 4-12% Bis-Tris plus (Thermo Fisher Scientific) and stained with the silver stain kit (Thermo Fisher Scientific) as per manufacturer’s instructions. The remaining samples were size-separated on a bolt 10 % Bis-Tris plus and stained with the Imperial protein stain (Thermo Fisher Scientific) according to manufacturer’s instructions. For each ChIRP experiments one gel slice from the SARS-CoV-2 and mock-infected samples were cut from the SDS-page. Gel slices from 5 independent replicates were sent for mass spectrometry analysis at Taplin Biological Mass Spectrometry Facility (Harvard Medical School) as previously described (Hafirassou et al., 2017).

### High-confidence RNA binding protein scoring

ChIRP-M/S data were analyzed using the online server (https://reprint-apms.org/) to run SAINTexpress (Teo et al., 2014). Total peptides counts from each of the five replicates, including both mock- and SARS-CoV-2-infected cells, were used as quantitative values to run the analysis. To determine the final list of high-confidence hits we applied stringent filtering criteria with MIST scoring > 0.7, and a SAINTexpress score above 0.78. Data shown in figure 1E correspond to the fold change scores (FC-A) calculated by SAINTexpress from the average mean of spectral counts.

### Identification of known and new interactors of coronaviruses in the ChIRP dataset

A recently published meta-analysis by Perrin-Cocon *et al*. was used as a reference for known interactions between coronavirus and host proteins (Perrin-Cocon et al., 2020). In this report, the literature was manually curated to assemble a dataset of 1,311 coronavirus-host interactions encompassing 1,140 distinct cellular proteins. Host proteins in the ChIRP dataset overlapping with those present in this reference interactome were considered as previously reported or « known » interactors of coronaviruses. They were displayed together with the interacting-coronavirus proteins as an interaction network using Cytoscape (Shannon et al., 2003). Cellular proteins in the ChIRP dataset that were not reported in Perrin-Cocon *et al*. were considered as « new » interactors of SARS-CoV-2 and coronaviruses in general. This list of new interactors was analyzed with the Functional Annotation Tool of the online knowledge base DAVID Bioinformatics Resources 6.8, NIAID/NIH (Dennis et al., 2003) to identify statistical enrichments in KEGG pathway annotations (Kanehisa and Goto, 2000).

### siRNA screen assay

An arrayed ON-TARGETplus SMARTpool siRNA library (Horizon Discovery) targeting 138 of the 142 RNA binding proteins significantly enriched in our ChRIP-M/S was purchased from Horizon Discovery. A549 cells stably expressing ACE2 were seeded in 24-well plate format (viral infection) or a 96-well plate (viability) and then were reverse transfected in duplicate with 30 nM of siRNA using the Lipofectamine RNAiMAX reagent (Invitrogen) according to manufacturer’s instruction. 48 hours posttransfection, cells seeded in a 24-well format were challenged with SARS-CoV-2 at a MOI of 0.05 in DMEM containing 5 % FBS. After three hours at 37°C, virus inoculum was removed and replace with fresh medium. 24 hours post-infection, cells and 200 μL of supernatants were collected for N protein staining and viral RNA quantification as described in ‘Viral infection quantification assays’. Throughout the screening assay, siRNA-transfected cells seeded in 96-well format were incubated for 72 hours at 37 °C and used to assess cell viability.

### Drug treatment and SARS-CoV-2 infection

Cells grown in 24-well plates were treated with the indicated compound concentrations for two hours prior infection and compounds were maintained throughout the course of infection. Cells treated with equivalent concentration of DMSO were used as control. The cells were challenged with SARS-CoV-2 at a MOI of 0.05 for 3 hours, then washed once with PBS and incubated with fresh media. 24 hours post-infection, cells and 200 μL of supernatants were collected for N protein staining and viral RNA quantification, as described in ‘Viral infection quantification assays’. In parallel cells seeded in 96-well format were incubated 24 hours with the drugs to assess cell viability

### Viral infection quantification assays

For infection quantification by flow cytometry analysis, 24 hours post-infection cells were trypsinized and fixed with 2% (v/v) paraformaldehyde (PFA) diluted in PBS for 15 min at room temperature. Cells were incubated for 45 min at 4°C with 1 μg/ml of the anti-N mAb diluted in permeabilization flow cytometry buffer (PBS supplemented with 5% FBS, 0.5% (w/v) saponin, 0.1% Sodium azide). After washing, cells were incubated with 1 μg/ml of Alexa Fluor 647-conjugated goat anti-mouse IgG diluted in permeabilization flow cytometry buffer for 30 min at 4°C. Acquisition was performed on an Attune NxT Flow Cytometer (Thermo Fisher Scientific) and analysis was done by using FlowJo software (Tree Star). To assess infectious viral particles release during infection by RT-qPCR, viruses were first inactivated by incubating the supernatants v/v with 1% Triton X-100 (Sigma) in PBS for 30 min under agitation at RT. Yields of viral RNA were quantified by real-time qPCR by using SARS-CoV-2 specific primers targeting the E gene with the Luna®Universal One-Step RT-qPCR Kit (New England Biolabs) in a LightCycler 480 thermocycler (Roche) according to the manufacturer’s protocol. The number of viral genomes is expressed as PFU equivalent/ml and was calculated by performing a standard curve with a similarly treated supernatant from a viral stock with a known titer as described by Gordon et al (Gordon et al., 2020b).

### Cell viability assay

Cell viability and proliferation were assessed using the CellTiter-Glo 2.0 Assay (Promega). Briefly, cells were plated in 96-well plates (5×10^3^). At specific times, 100 μl of CellTiter-Glo reagent mixed v/v with PBS were added to each well. After 10 min incubation, 90 μl from each well were transferred to an opaque 96-well plate (Cellstar, Greiner bio-one) and luminescence was measured on a TriStar2 LB 942 (Berthold) with 0.1 second integration time.

## QUANTIFICATION AND STATISTICAL ANALYSIS

### Statistical analyses

Graphical representation and statistical analyses were performed using Prism7 software (GraphPad Software). Unless otherwise stated, results are shown as means +/- standard deviation (SD) from at least 2 independent experiments in duplicates. Differences were tested for statistical significance using One-way or Two-way Anova with multiple comparisons post-test.

Protein functional enrichment was performed in Cytoscape (Version. 3.8.2). Gene Ontology (GO) Biological process were enriched with the ClueGO (Bindea et al., 2009) extension (Version. 2.5.7) using a two-sided hypergeometric test with Bonferroni correction. Adjusted p-values < 0.5 were considered as significant. P-values and percentage of represented genes per GO terms were computed to select the most enriched functional families. This was illustrated with ggplot2 package (Version. 3.3.2) in R Studio interface (Version. 1.2.1335). Most representative terms were represented in networks using the Cytoscape’s Genemania extension (Version. 3.5.2). Comparison of functional enrichment between our datasets and previous studies was performed in Metascape (Zhou et al., 2019b). GO Biological process with p-value < 0.01, enrichment score > 1.5 and overlap = 4 were selected.

## Notes

### Competing Interest Statement

The authors have declared no competing interest.

